# Conjugative Transfer from *Escherichia coli* to Gram-positive Bacteria: A Systematic Review and Meta-Analysis

**DOI:** 10.1101/2025.03.11.642732

**Authors:** Supanida Chuenaem, Chanon Jaichuen, Sopida Wongwas, Pakpoom Subsoontorn

## Abstract

Conjugative DNA transfer is a major driver of microbial evolution and an essential tool for biotechnological applications. While conjugation between Gram-negative and Gram-positive bacteria has been observed, its efficiency and underlying principles remain poorly understood. This systematic review and meta-analysis assess the success and influencing factors of *Escherichia coli*-to-Gram-positive conjugation. A systematic search of the PubMed, Crossref and Web of Science database (up to July 2025) identified 41 studies reporting *E. coli*-to-Gram-positive conjugation, comprising 645 measurements. Studies were included based on experimental evidence of conjugative transfer and reported efficiency values. Data extraction was performed manually, and statistical analyses were conducted to identify key trends. We found that *E. coli*-to-Gram-positive conjugation is significantly less efficient than *E. coli*-to-*E. coli* or *E. coli*-to-other Gram-negative bacteria. However, certain recipient strains and optimized conditions enable surprisingly high efficiencies, within 1–2 orders of magnitude of *E. coli*-to-*E. coli* conjugation. Strategies that improve efficiency include altering plasmid methylation patterns, optimizing cation concentrations, and modifying donor-recipient ratios. Limitations include potential bias toward well-studied bacterial groups (e.g., *Streptomycetaceae*), variability in experimental conditions, and incomplete reporting in some studies. Further research should explore additional recipient strains, refine conjugation mechanisms, and optimize transfer conditions to enhance DNA delivery to non-model microbes. Understanding these processes may pave the way for more efficient and universal DNA transfer methods across diverse microbial taxa.

**Importance:** Bacteria have an incredible ability to share genetic material with each other, a process that drives evolution and enables survival in changing environments. Conjugation allows bacteria to pass DNA from one cell to another. Scientists have used this process to engineer microbes for medicine, agriculture, and environmental cleanup. However, transferring DNA between very different types of bacteria, especially from *E. coli* (a Gram-negative bacterium) to Gram-positive bacteria, has remained a challenge. This study reviewed over 600 experiments to understand when and how such transfers work best. Surprisingly, the researchers found that under certain conditions, DNA can move across this divide much more efficiently than previously thought. These insights could help scientists develop universal DNA delivery tools, unlocking the potential of hard-to-engineer bacteria for biotechnology. In a world that increasingly relies on microbes for sustainable solutions, improving how we “send instructions” into these microbial workers is a critical step forward.

## Introduction

Conjugative transfer could mobilize DNA across extremely diverse host cells. Previous studies reported the uses of broad-host-range conjugative transfer machinery, such as those from RP4 and Ti plasmid, for delivering DNA to various gram-negative and gram-positive bacteria as well as eukaryotes ranging from yeast to diatom, algae and even mammalian cells [1-6]. This capability implies pivotal roles of conjugative plasmids as hyper promiscuous agents for genetic exchanges in microbial evolution. For applications in synthetic biology, conjugative transfer is among primary tools for genetic manipulation. This mechanism allows for the delivery of a long (up to millions of nucleotides) DNA [7] and allows for delivery to recipient cells that might be immune to simpler methods such as direct DNA transformation [8]. Efficient and broad-spectrum DNA delivery can be used for engineering novel hosts of microbial cells factories and for spreading desirable genetic features (such as pollution degrading enzymes, CRISPR/Cas targeting antibiotic resistant genes, etc.) *in situ* [9-13].

What is the actual range of recipient cells for conjugative transfer? What are factors determining this range and DNA transfer efficiency into different recipients? Is there any general principle for achieving efficient transfers regardless of recipient cell types? Answers to these questions not only help us explain and predict the impact of conjugative transfer in microbial evolution but also suggest possible strategies toward developing universal DNA delivery tools for undomesticated microbes. While it is impossible to experimentally test conjugative transfer to all known recipient cells, previous studies offer insights related to these questions. One approach is an *in situ* conjugation experiment, where conjugative or mobilizable plasmids of interest are delivered to microbiota in soil, sludge, gut and skin [3, 14-16]. Then, transconjugants are recovered (using fluorescent activated cell sorters or antibiotic selection) and taxonomically classified (using metagenomic sequencing of markers such as 16s rRNA genes). In a single experiment, this approach can give an overview of recipient ranges as well as relative transfer efficiency among many possible recipients in the microbiota. However, factors such as abundance and distribution of cell in microbiota as well as abiotic conditions (such as chemical ions, humidity, granularity of substrate, etc.) can confound the analysis. Another complementary approach is to separately measure conjugation efficiencies for each individual recipient cell types. This way, experimenters can have better control of conjugation environment and directly measure absolute efficiency of transfer. Still, this approach is laborious so each individual study usually measures conjugation efficiency in few recipient cell types and explores few factors related to conjugation efficiency.

Systematic review and meta-analysis pool data from multiple studies to draw conclusion beyond what individual study can offer. This approach allows us to systematically search literature, curate content, extract data and perform analysis with minimal bias toward certain hypothesis. It helps us assess state-of-the-art as well as identifying knowledge gaps and discrepancies in previous studies. While systematic review and meta-analysis is common in clinical research, this approach has rarely been applied in basic science. In the context of studying plasmid conjugation, three seminal studies demonstrated how this approach can help assess contributions of various factors on conjugation efficiency. Pooling data from 34 studies with a total of 128 measurements, Hunter et al (2008) identified types of donor cells, recipient cells, transferred plasmid and conjugation setup (*in vitro* vs *in vivo*) as main contributors [8]. The authors also built general linear models, using these factors as input variables, that can decently predict conjugation efficiency. Pooling data from 33 studies with a total of 612 measurements, Sheppard et al (2020) additionally identified plasmid repression as a major contributor of conjugation efficiency [17]. Lastly, using data from 32 studies a with a total of 583 measurements, Alderliesten et al (2020) concluded that further taxonomic distance between donor and recipient cells is associated with lower conjugation efficiency [18]. This association, however, is only significant when conjugation happened in liquid media and not on solid agar surface.

Unfortunately, these systematic review and meta-analysis works barely offer insight about recipient ranges for conjugative DNA transfer and key factors affecting such ranges. The limitation stems from how these systematic reviews were designed. First, the collected data tend to bias toward conjugative transfer between closely related cell types. For example, in the study by Hunter et al (2008) over 80% of measurements were from conjugation between bacteria in the same gram [8]. Alderliesten et al (2020) focused on measurement of conjugation from *Escherichia coli* donor and diverse recipients. Still, only 2 out of 583 measurements have gram positive bacteria as recipient [18]. Second, these systematic reviews did not further investigate the extent to which different factors, both genetic and environment, affect conjugative DNA transfer to an individual recipient species or strain. Rather, these systematic reviews pooled together measurements from various recipient cell types; some of these recipient types may take up DNA much more or much less efficient than others under certain circumstances. This would severely skew the overall conjugation efficiency measures of the pooled data. Third, all these systematic reviews only use data from self-transmissible conjugative plasmids but exclude data from mobilizable plasmids (which can also be conjugatively transferred using conjugation machinery provided in trans from another plasmid or genome). Many mobilizable plasmids developed for DNA delivery applications have origin of replications suitable for recipient cells or can be integrated into recipient genomes. While these plasmids are not ‘natural’, they can still provide valuable information about recipient ranges for conjugative DNA transfers. They allowed us to measured DNA transfer range and efficiency without being confounded by an ability to maintain the DNA in recipients.

Here, our systematic review and meta-analysis address limitations above while focus on conjugation from *E. coli* to gram-positive bacteria. Not only is this type of conjugation severely underrepresented in previous systematic reviews but gram-positive bacteria are also an important DNA delivery target in various biotechnological applications. Gram-positive bacteria such as some *Bacillus* and *Lactobacillus* are probiotics for human and animals as well as growth promoting and biocontrol agents for plants [19-21]. *Streptomyces* have been used for production of antibiotics and other valuable secondary metabolites [22]. *Staphylococcus* and *Streptococcus* are among pathogenic bacteria with great public health concerns. Many strains of these gram-positive bacteria are difficult to DNA transform. Thus, conjugation remains the primary methods for DNA delivery, either for strain improvement or genetic manipulation in basic research. Additionally, conjugation between gram-negative (*E.coli*) to gram-positive could offer insight about astonishing promiscuity of this DNA transfer mechanism. Gram-negative and gram-positive are evolutionarily distant; their external structures (cell walls and outer membranes) are drastically different. There is unlikely any evolutionary force selecting for gram-negative to gram-positive conjugation. Nor should there be recipient specific receptors or molecular interaction facilitating this process. Still, mere existence of gram-negative to gram-positive conjugation implies universality of this DNA transfer process. By understanding the limits of this types of conjugation and factors influencing its efficiency, we may take the first step toward developing universal DNA delivery platform that can cross any taxonomical distance and structural diversity. This systematic review and meta-analysis aim to assess the efficiency of *E. coli*-to-Gram-positive conjugation, identify key influencing factors, and determine general principles that may enhance inter-gram conjugation efficiency.

## Results and Discussion

### Identification of relevant studies

Our search found 12,821 potentially relevant studies (Fig 1). 10,393 studies were excluded as their titles were not related to conjugation between *E.coli* and Gram-positive bacteria. The remaining 275 studies were further evaluated based on their abstracts. Studies that did not explicitly mention conjugation between *E.coli* and Gram-positive bacteria were excluded, resulting in 132 studies remaining. After screening full text, we excluded 94 studies and have 38 studies. In addition, we included three relevant studies that were not retrieved by our search strategy but were previously known to our research team. This brought the total to 41 studies for data extraction. The data extracted from these studies encompassed a total of 740 measurements, which formed the basis for our comprehensive analysis and findings in this systematic review. Note that, only 645 measurements are from *E.coli*-to-gram-positive conjugation; the rest are from *E.coli*-to-*E.coli* or gram negative conjugation, which we also use for comparison.

**Fig. 1.**
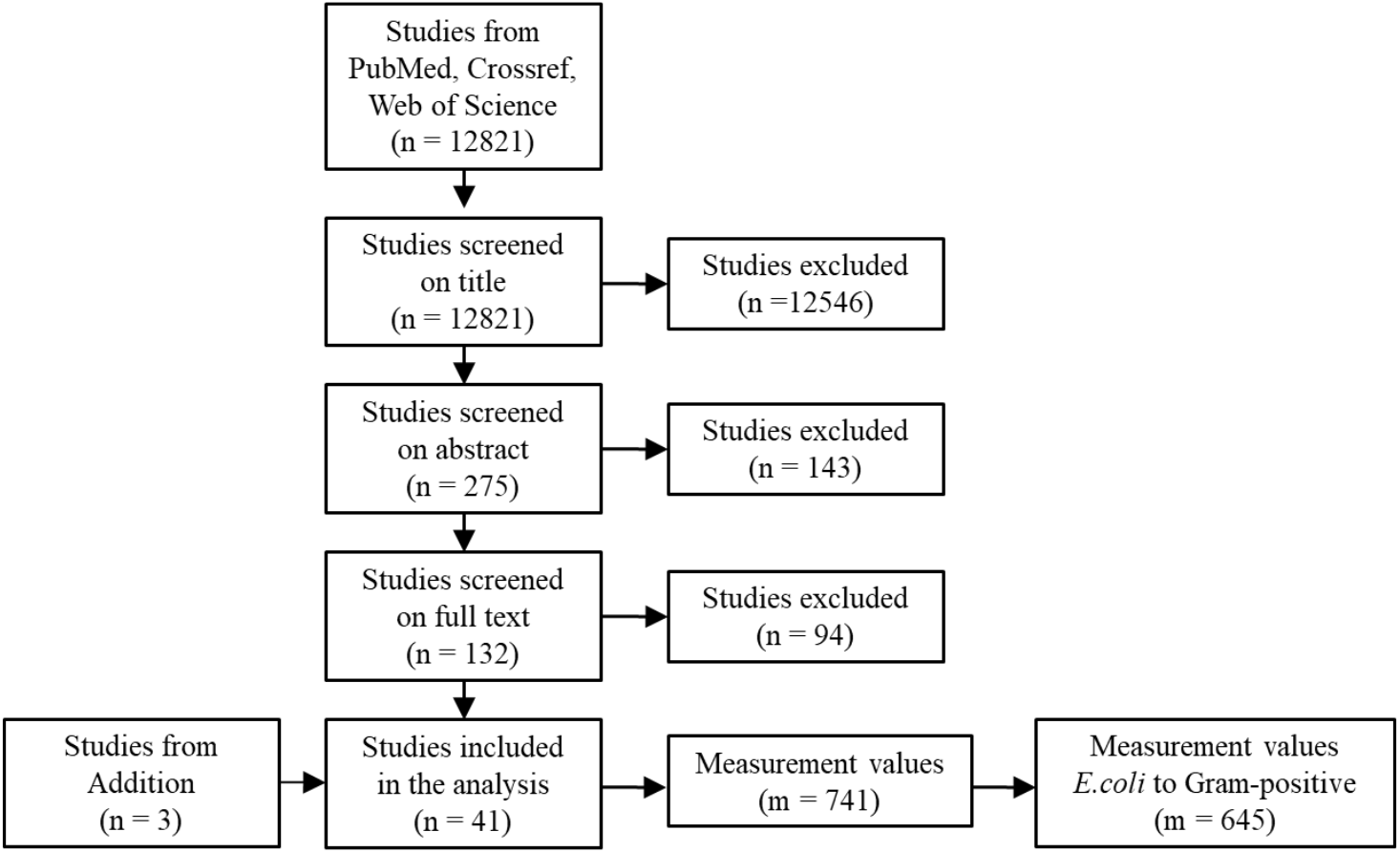
Flowchart depicting the search results and the selection of studies. n: number of studies, m: number of measurements

Some studies reported multiple measurement values of conjugation efficiency from the same set of donor, recipient and transfer plasmid because the authors attempted to optimize conjugation environment. For example, Wang et al (2014) varied the concentration of cation and media types to find optimal condition for *E.coli* to *Streptomyces* conjugations [6]. Consequently, the authors reported up to 41 measurement values of conjugation efficient from the same set of donor, recipient and transferred plasmid. When we tried to answer the question “How efficient can conjugation efficiency from *E.coli* to bacteria X be?”, we should consider measurement taken under the most optimal condition. Otherwise, measurement data from suboptimal conditions would obscure our analysis. Thus, the 645 measurement values of *E.coli* to gram-positive conjugation were reduced to 380 measurement values, each of which was taken from a unique combination of donor, recipient, transferred plasmid and study. We will use these 380 measurement values for subsequent analysis in Fig 2 and Fig 3.

**Fig. 2.**
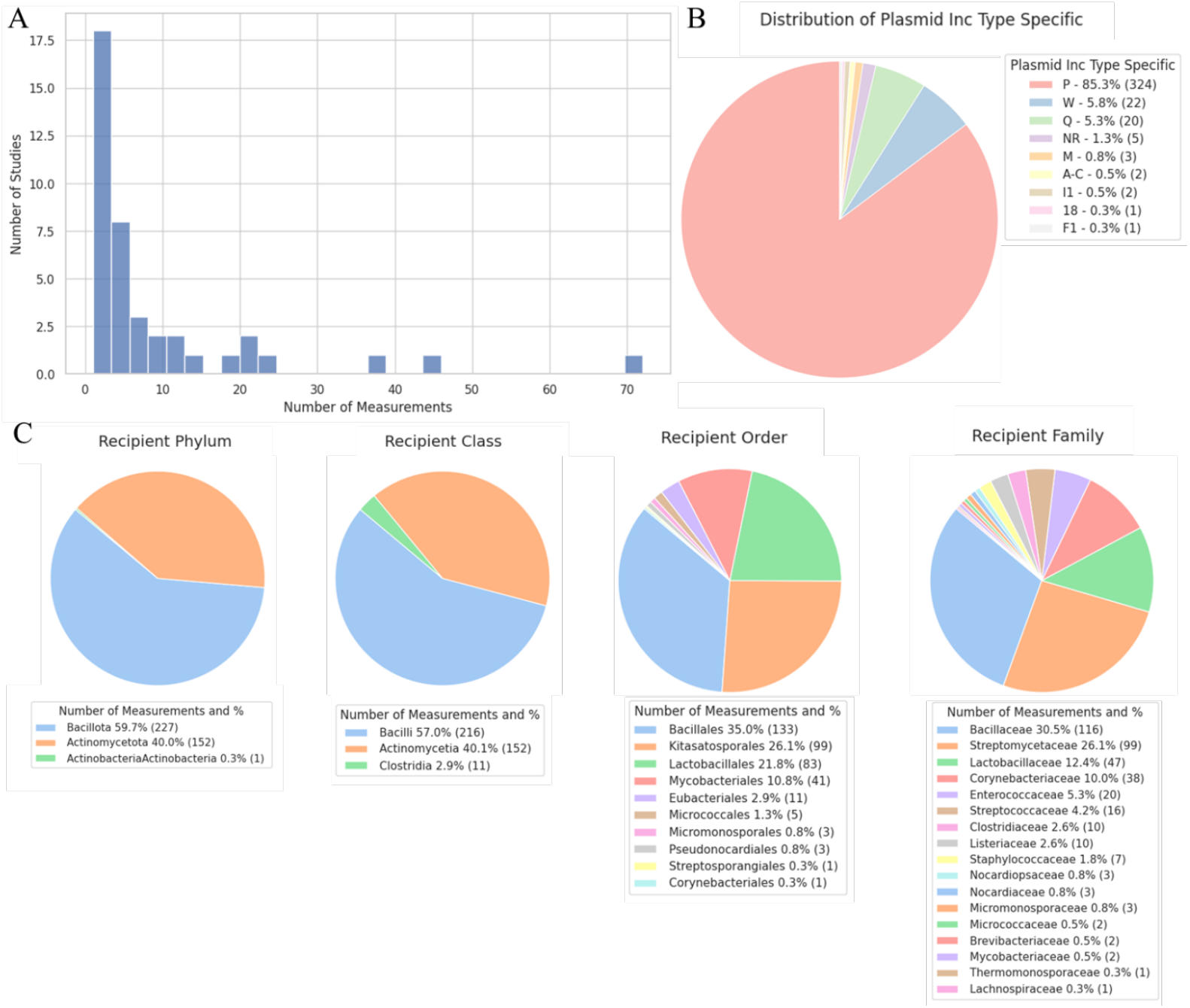
Distribution of measurement numbers and types. (A) histogram showing the distribution of numbers of measurements from selected studies. (B)-(C) pie charts showing the distribution of conjugation mechanisms and recipient classification from all measurements.

**Fig. 3.**
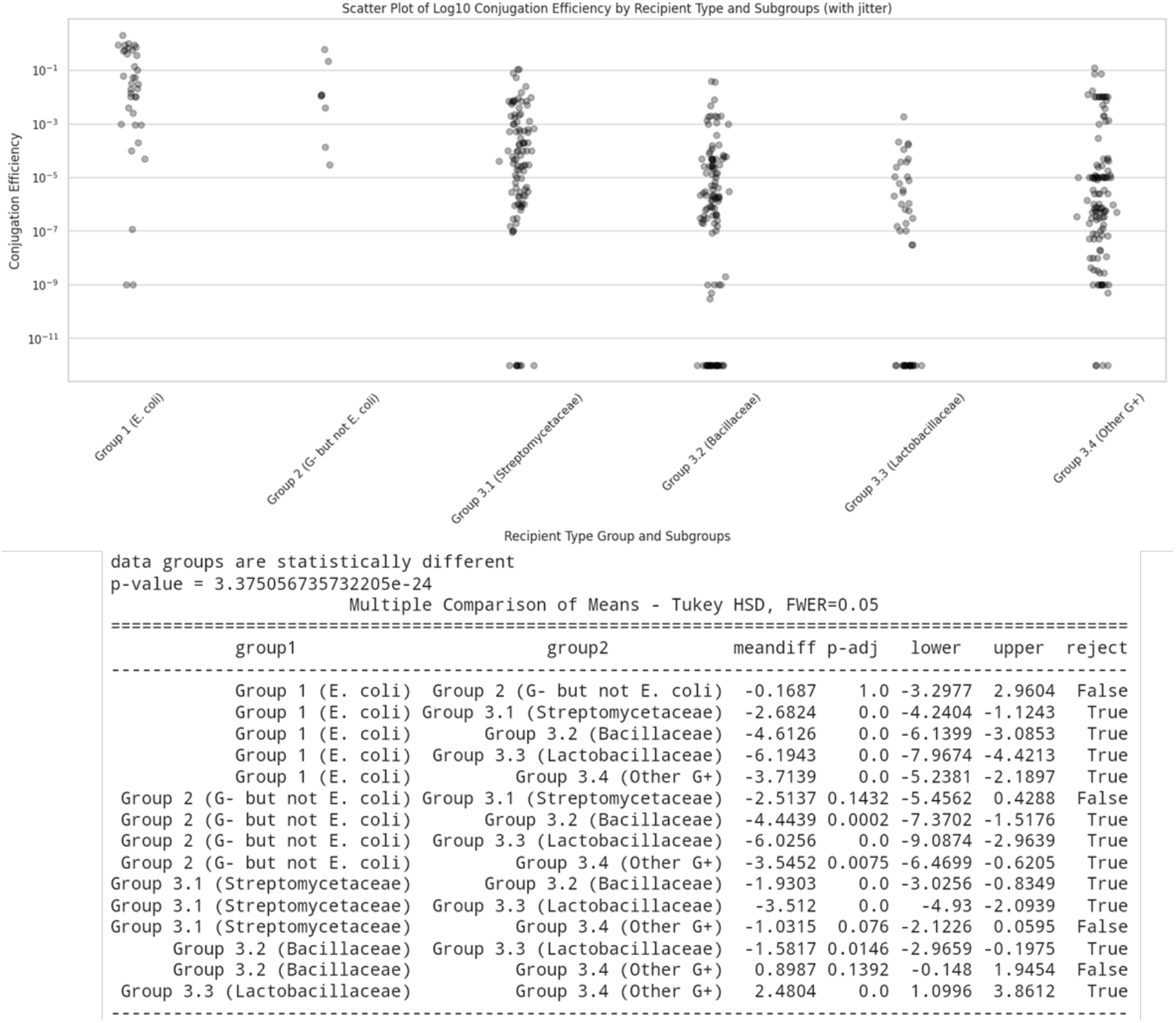
Conjugation efficiency from *E.coli* donor to various groups of recipient bacteria. Efficiencies were measured as transconjugant-per-recipient. Conjugation efficiencies below detection limit were set to 1E-12 in this plot and statistical analysis.

### Overall trends of selected studies

Only ten studies provided more than 10 measurement values (Fig 2A). Together, measurement values from these ten studies accounts for almost one fourth of all measurement values in our meta-analysis. Conjugation machinery from plasmid incompatibility group P (IncP) was the most frequently used, accounting for 85% of all measurement values (Fig 2B). Regarding recipient cell types, all recipients used in these measurements either belong to *Actinomycetota* or *Bacillota* phylum (Fig 2C). *Bacillota* group consists of various taxa at class, order and family level while *Actinomycetota* group is almost completely dominated by *Streptomycetaceae*.

### Conjugation Efficiencies from *E.coli* to Various Recipient Groups

We compared reported conjugation efficiencies from *E.coli* to six different bacteria groups: 1) *E.coli*, 2) other gram-negative bacteria, 3.1) *Streptomycetaceae*, 3.2) *Bacillaceae,3.3)Lactobacillaceae*, and 3.4) other gram-positive bacteria (Fig 3). All data points for group 3.1-3.4 came from the 380 measurement values above. Data points for group 1-2 came from selected studies that also reported conjugation efficiencies from *E.coli* to gram-negative bacteria [5, 18, 23-26].

Statistical test showed that mean *E.coli*-to-*E.coli* conjugation efficiency is over two orders of magnitude higher than mean conjugation efficiency of *E.coli* to all four groups of gram-positive bacteria. According to this data, mean conjugation efficiency from *E.coli* to other gram-negative bacteria is not significantly different from *E.coli*-to-*E.coli* conjugation. Among four groups of gram-positive bacteria, conjugations with *Streptomycetaceae* as recipient have highest mean conjugation efficiency followed by other gram-positive and *Bacillaceae* and finally *Lactobacillaceae* with the lowest mean efficiencies. Difference in conjugation efficiencies among groups of gram-positive recipients may not be conclusive, due to bias from conjugation machineries uses and differences in setup across studies. For example, when we use only data from studies that have *E.coli*-to-*E.coli* conjugation as measurement controls or when we only measurement from experiment with Inc P conjugation machineries, some of these differences in conjugation efficiencies among groups of gram-positive recipient disappeared (Table S1, Fig S1 – S3). Nonetheless, the observed difference in conjugation efficiencies between *E.coli* recipient and gram-positive recipient persist across different choices of data set (Table S1).

Our result agrees with the observation of Hunter et al (2008) that gram-negative to gram-negative conjugation is more efficient than gram-negative to gram-positive conjugation [8]. However, Alderliesten et al (2020) reported that taxonomic distances between donor and recipient were not associated with conjugation efficiency on solid media [18]. In our case, nearly all conjugation experiments were performed on solid media. *E.coli* to *E.coli* and *E.coli* to other gram-negative conjugations in our case definitely have smaller taxonomic distance than *E.coli* to gram-positive conjugation. At glance, there seems to be discrepancy between our observation here and Alderliesten et al (2020) conclusion. A possible explanation is that Alderliesten et al (2020) only have a single measurement of *E.coli* to gram-positive conjugation on solid media, not enough data to see significant difference in statistical test.

We notice that maximum conjugation efficiency for each gram-positive group was surprisingly high, reaching 1E-1 for *Streptomycetaceae* and *Bacillaceae* and reaching 1E-3 for *Lactobacillaceae* and other gram-positive. Comparing conjugation efficiencies across studies could be challenging as each study usually has different experiment setup. Fortunately, among our selected studies, six reported both *E.coli* to *E.coli* and *E.coli*-to-gram-positive conjugation, allowing for direct comparison within the same study. Three studies reported that *E.coli* to gram positive (*Streptomyces, Bacillus, Lactobacillus*) conjugation efficiency could be within 1-2 orders of magnitude from *E.coli*-to-*E.coli* conjugation efficiency [20, 22, 23]. These results imply that it is theoretically possible to achieve efficient conjugation from gram-negative to gram-positive bacteria, despite their taxonomical distance and striking differences in their cell wall and membrane structures. Generally observed low efficiencies of conjugation between gram-negative and gram-positive could result from suboptimal conjugation environment or strain-specific defenses against foreign DNA in recipient cells. The overall certainty of evidence in this systematic review is influenced by variability in experimental conditions and reporting standards. While sensitivity analyses (Table S1) confirm the robustness of major trends, differences in recipient strains, conjugation machineries, and measurement techniques contribute to uncertainty. No formal certainty assessment framework (e.g., GRADE) was applied, but confidence in findings is supported by consistency across independent studies.

Given observed differences in conjugation efficiencies among four groups of gram-positive recipients, we asked whether such difference results from recipient taxa or any other factors relate to experimental designs. Strain is the lowest level of taxa. Still, even for the same recipient species, conjugation efficiencies could differ by several orders of magnitude across different recipient strains. Thus, we further expand our visualization of conjugation efficiencies to include information about recipient strains and other independent variables (Fig 4). Here, we started with 645 measurement values of *E.coli* to gram-positive conjugation. We chose to display results from recipient strains that have at least two data point (i.e., the authors have tried to vary at least one independent variable related to conjugative transfers to this recipient strain). This would allow us to explore the effects of environmental variables (e.g. media, temperature, etc.) as well as genetic variables (donor strain and plasmids). These variables influencing conjugation efficiencies could be used for optimizing conjugation (Table 1)

**Fig. 4.**
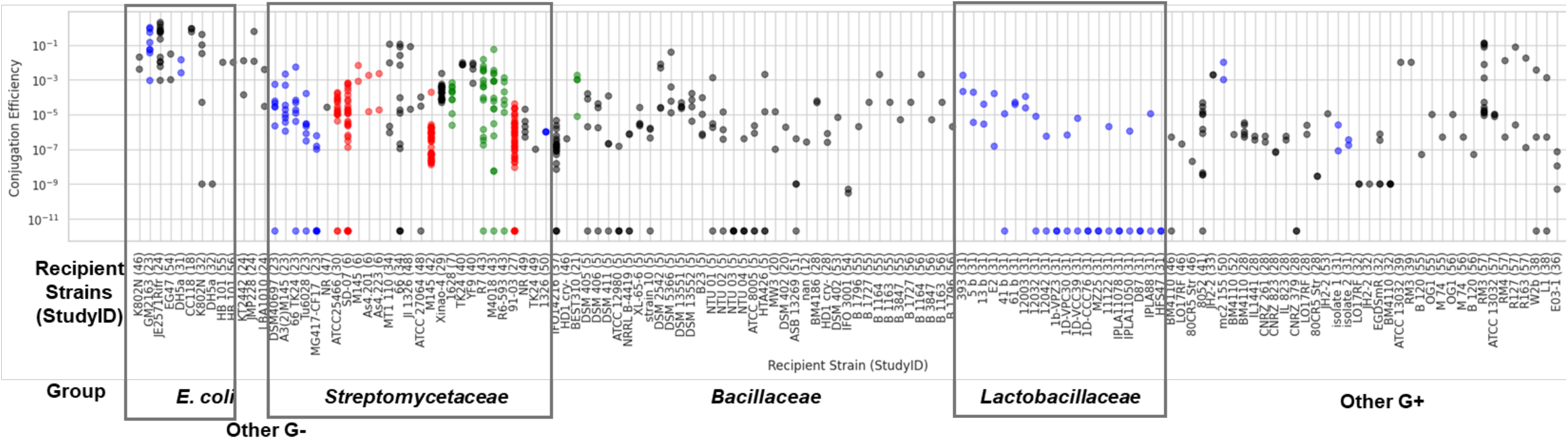
Conjugation efficiency from *E.coli* donor to various recipient strains. Colors of data points indicate types of independent variables. Black: donor strains and transferred plasmids; blue: conjugation machineries; red: media formulation; green: time, temperature or cell density.

**Table 1.** Strategies for improving conjugation efficiency.

| Strategies | Recipient | Fold Improvement <sup>[StudyID]</sup> |
| --- | --- | --- |
| Change methylation pattern | *Bacillus* | 2.0E+9<sup>[5]*</sup> |
|  | *Streptomyces* | 1.4E+1<sup>[40]</sup>, 5.5E-1<sup>[[27]</sup>, 1.1E+11<sup>[34]*</sup> |
| Change conjugation machinery | *Streptomyces* | 5.3E+9<sup>[23]*</sup> |
|  | *Lactobacilli* | 1.1E+7<sup>[31]*</sup> |
|  | *Staphylococcus* | 3.1E+1<sup>[31]</sup> |
| Change machinery location | *Streptomyces* | 4.7E+7<sup>[41]*</sup> |
| Add Al<sub>2</sub>O<sub>3</sub> nanoparticle | *Streptomyces* | 1.5E+4<sup>[42]</sup> |
| Add cations (Mg<sup>2+</sup>, Ca<sup>2+</sup>) | *Streptomyces* | 1.8E+6<sup>[27]*</sup>, 6.4E+8<sup>[6]*</sup>, 4.7E+0<sup>[29]</sup>, |
| Change donor and recipient ratio | *Streptomyces* | 4.2E+1<sup>[27]</sup>, 3.5E+1<sup>[29]</sup>, 1.1E+1<sup>[30]</sup> |
|  | *Enterococcus* | 1.1E+2<sup>[33]</sup> |
| Heat shock temperature before conjugation | *Streptomyces* | 4.4E+0<sup>[43]</sup>, 7.1E+0<sup>[29]</sup> |
|  | *Corynebacterium* | 1E+2<sup>[39]</sup> |
| Incubate time after heat shock | *Streptomyces* | 9.2E+0<sup>[9]</sup> |
| Optimize recipient culture time | *Streptomyces* | 2.5E+9<sup>[43]*</sup> |
| Adjust incubation time | *Streptomyces* | 2.2E+0<sup>[29]</sup> |

Fold Improvement: maximal reported fold improvement among all tested strains in each study.
\*Conjugation efficiency before improvement is below detection limit reported in the study.

Among four groups of gram-positive recipients, we can clearly see the diversity of independent variables explored on *Streptomycetaceae* conjugation (diversity of data point colors in Fig 4). Thus, it is more likely that optimal conditions for conjugative transfer have been found in this recipient groups compared to other gram-positive groups. This could explain why mean conjugation frequency of this group appears to be the highest (Fig 3, Table S1). Environment conditions could have huge effects. For example, in some studies of *Streptomyces*, varying levels of cation (particularly, Ca^2+^ and Mg^2+^) and other media compositions cause conjugation efficiencies to change by several orders of magnitude [27, 28]. This observation raise question on validity of previous systematic reviews and meta-analyses. While these previous works did explore some environmental variable such as media richness, temperature, mating pore size, etc., they have not looked closely into whether these variables were applied differently in different recipient groups. As in our case, almost all measurement data related to environmental variables could come from one group (i.e. *Streptomyces*). If such data (green and red data points in Fig 4) was all included in the meta-analysis (Fig 3), they would bring down the mean conjugation efficiency (as many of these data points were conjugation performed under suboptimal condition). On the contrary, if only the highest conjugation efficiency measures were selected for each set of donors, recipient and transferred plasmid (as we did in this meta-analysis), they would be likely to bring up mean conjugation efficiencies of the group (as conjugation were performed in the most optimized condition).

Wang et al. (2014), Sun et al. (2014), and Du et al. (2012) collectively examined the influence of MgCl_2_ concentration on bacterial conjugation efficiency [6, 29, 30]. Sun et al (2014) and Du et al (2012) [17, 30] both observed that a MgCl_2_ concentration range of 20–40 mM resulted in optimal conjugation efficiency, while Wang [6] reported a higher optimal concentration at 60 mM. The variations in optimal MgCl_2_ concentrations across studies may be attributed to differences in the recipient strains used. The underlying mechanisms by which MgCl_2_ enhances conjugation efficiency remain unknown. However, we hypothesize that MgCl_2_ may influence cell membrane stability and plasmid interactions, facilitating plasmid uptake into recipient cells and leading to improved conjugation efficiency [10]. Further investigations are needed to elucidate the precise mechanisms by which MgCl_2_ contributes to conjugation efficiency. Future studies should focus on experimentally determining whether MgCl_2_ affects membrane permeability, pili formation, or other conjugation related cellular processes. Additionally, evaluating the role of MgCl_2_ across different bacterial species and genetic backgrounds may provide further insights into its effects on plasmid transfer.

Conjugation efficiencies data of *Lactobacillaceae* and *Bacillaceae* came almost exclusively from only two studies [26, 29]. In the study by Samperio et al (2021), various *Lactobacillaceae* strains were conjugated with *E.coli* donor carrying conjugation machinery from either RP4 plasmid (from IncP) or R388 plasmid (from IncW) [31]. For every tested *Lactobacillaceae* strain, conjugative transfer by RP4 is far more efficiency than by R388. In fact, R388 can only deliver a plasmid to 6 out of 20 tested strains while RP4 can delivered to 14 strains. When the conjugation efficiency data from RP4 and R388 delivery were pooled together in the meta-analysis (Fig 3), it was not surprising that mean conjugation efficiency was low. Other selected studies in our systematic review reported similar trend: RP4 system is generally more efficient than other alternative conjugation machinery (Fig S2). The only exception is in two *Staphylococcus epidermidis* recipient strains where Samperio et al found R388 system to be slightly more efficient than RP4. Most of our selected studies used only RP4 system and thus mean conjugation efficiencies were not lowered by data from other less efficiency conjugation systems. Again, this kind of bias have not been discussed in previous meta-analysis works. It is possible that reported mean conjugation efficiency for certain groups of recipients appear to be less simply because they have been tested on less efficient conjugation systems.

Interestingly, two studies have reported natural plasmids capable of transferring and replicating in both Gram-positive and Gram-negative bacteria [32, 33]. These include the conjugative plasmid pIP501, originally isolated from Enterococcus faecalis, and the mobilizable plasmid pIP823, from Listeria monocytogenes. Unlike systems described in other studies, these plasmids can self-transfer or be mobilized in both directions from Gram-negative to Gram-positive bacteria and vice versa. This is particularly surprising given the substantial differences in cell wall composition and outer membrane structures between these two groups, which are typically thought to prevent the same conjugative pili and DNA transfer machinery from functioning in both direction. These findings highlight that the Gram classification of a recipient bacterium does not necessarily constrain plasmid transfer. Moreover, they suggest that naturally occurring plasmids with broad-host-range transfer capabilities may play a critical role in promoting horizontal gene transfer, including the spread of antibiotic resistance, across diverse microbial communities.

The study by Tominaga et al (2016) provides 72 data points of conjugation efficiency measurements, derived from testing 18 *Bacillus* or *Geobacillus* recipient strains with four different *E. coli* donor strains [5]. These donor strains were engineered to imprint distinct DNA methylation profiles on the transferred plasmid, with the rationale that mimicking the recipient’s native methylation patterns would allow the incoming DNA to evade restriction cleavage by host restriction-modification (R-M) systems. The study reported striking variations in conjugation efficiencies, spanning several orders of magnitude depending on the donor strain used. This highlights the critical role of DNA methylation in determining the success of conjugative plasmid transfer from *E. coli* to gram-positive bacteria.

Research on conjugation into *Streptomyces* species has focused on overcoming methylation barriers using unmethylated donor DNA. This is achieved by using the methylation-deficient *E. coli* strain *ET12567*, which lacks both Dam (DNA adenine methylase) and Dcm (DNA cytosine methylase) activities (dam-/dcm-). When this non-methylating strain is combined with a conjugative helper plasmid like pUB307, it has demonstrated a 10^4^-fold increase in conjugation efficiency when transferring DNA to *Streptomyces coelicolor* and *Streptomyces lividans* compared to standard methylation-competent donor strains [34]. The success of this approach lies in the strain genetics (*ET12567*’s dam-/dcm-status) rather than any methylation characteristics of the helper plasmid pUB307 itself. This strain-plasmid combination allows DNA to be transferred without the methylation patterns that would typically trigger restriction systems in the *Streptomyces* recipient.

Another key advancement in *Streptomyces* conjugation has been the development of *ET12567*(pUZ8002). Recent research by Larcombe et al (2024) has revealed that pUZ8002, a helper plasmid, originated from the Spanish *P. aeruginosa* plasmid pUZ8, rather than RP4 as previously thought [35]. While pUZ8002 contains the Tra and Trb genes essential for conjugation, it features an engineered inactive oriT created through four specific nucleotide modifications. This modification eliminates the TraI-binding and nic site, preventing self-conjugation while maintaining the ability to mobilize other plasmids efficiently. This characteristic has made pUZ8002 a cornerstone tool in *Streptomyces* molecular genetics [36-38].

These collective findings emphasize that methylation profiles strongly influence conjugation efficiency between gram-negative and gram-positive bacteria, primarily due to the latter’s stringent restriction-modification systems. Whether through the removal of methylation (dam-/dcm-donors) or by mimicking recipient methylation patterns, researchers have developed effective strategies to enhance plasmid transfer success. This work demonstrates that overcoming methylation pattern mismatches between donors and recipients is crucial for successful intergeneric conjugation, particularly when working with gram-positive bacteria and their complex protective mechanisms.

One of the most notable enhancements in conjugation efficiency from *E. coli* to gram-positive recipients was observed in *Corynebacterium glutamicum* following heat treatment of the recipient cells [39]. Under standard conditions, transfer frequencies of the mobilizable shuttle vector pECM1 from *E. coli S17-1* to *C. glutamicum* were relatively low, ranging between 10^−6^ and 10^−7^ per donor. However, when recipient cells were subjected to a 9-minute heat treatment at 48.5°C prior to mating, conjugation frequencies markedly increased to 10^−2^– 10^−3^ per donor, representing a 1,000-to 10,000-fold enhancement. This dramatic improvement was attributed to the inactivation of recipient restriction systems, which otherwise degrade incoming foreign DNA. Indeed, when a restriction-deficient (Res^−^) mutant of *C. glutamicum* was used, similar transfer frequencies were observed even without heat treatment, confirming the role of restriction barriers in limiting conjugation. Moreover, no additional increase was conferred by combining the Res^−^ background with heat treatment, supporting the notion that heat exposure primarily acts to suppress restriction activity. When applied across multiple wild-type coryneform strains, heat treatment consistently elevated transfer frequencies, although efficiency varied by taxonomic group. All group A strains and most group B strains showed detectable transfer, whereas group D strains remained refractory. This indicates that while heat treatment is a powerful enhancer of DNA uptake, intrinsic compatibility factors such as replicon compatibility and cell envelope properties may still govern ultimate success. Taken together, these findings demonstrate that heat pretreatment of recipient cells can be a simple yet effective strategy to overcome interspecies conjugation barriers primarily by transiently impairing recipient’s restriction systems. Such methods may facilitate broader applications of conjugative DNA delivery into gram-positive bacteria, especially those with traditionally low transformation and conjugation efficiencies.

### Conclusion and Future Directions

To the best of our knowledge, our systematic review and meta-analysis is the first to explore interspecies conjugation with emphasis on *E.coli* to gram-positive bacteria conjugation. In attempt to answer the key questioned posted in the introduction on 1) range of conjugative transfers, 2) factors influencing conjugation efficiency, 3) general principle for achieving efficient transfers, we systematically searched, screened, extracted and analyzed data about *E.coli* to gram-positive conjugation from 41 studies encompassing over 600 measurement values of conjugation efficiencies. Unlike previous systematic review and meta-analysis works, we not only pooled data to perform statistics but also dive deep into detail experimental results from each individual selected study. From this exercise, we draw the following lessons and propose the following suggestions.

1. We have assembled the most comprehensive list of gram-positive bacteria that can received plasmids from *E.coli* via conjugative transfer. This list consists of 19 gram-positive families and 31 genera where directed measurement data of conjugation efficiencies is available in literatures. The true conjugative transfer ranges in gram-positive is likely to be much larger than this. Future works could involve expanding the initial search term to include more gram-positive bacteria and incorporate data from *in situ* conjugative transfer studies (Fig S4).
2. Given their taxonomical distance and structural difference, it is intuitively believed that *E.coli* to gram-positive should be much less efficient than *E.coli* to *E.coli* conjugation. However, the highest reported conjugation efficiencies to *Streptomycetaceae, Bacillaceae* and *Lactobacillaceae* family can be within 1-2 order of magnitude from conjugation efficiency to *E.coli.* Thus, under a well optimized conjugation environment and in the absence of strain-specific defense against foreign DNA, efficient conjugation across such vast taxonomical distance and structural difference is possible. This observation has important implication in the development of a universal DNA delivery platform. Future works should first confirm all these observations of efficient *E.coli* to gram-positive conjugation (which come from many studies spanning many decades) in the same experimental setting and then dive into their detail mechanisms.
3. Conjugative transfer to certain gram-positive taxa, particularly *Streptomycetaceae*, were studied more extensively than others possibly due to their industrial significance and the lack of other efficient DNA delivery approach. Numerous strategies have been developed to boost conjugation efficiency ranging from optimizing media, temperature and incubation time to designing specific donor strains and transferred plasmids. It is thus not surprising that the highest conjugation efficiency reported for this group reached up to over 1E-1, comparable to that of *E.coli* to *E.coli* conjugation. Future works should attempt to apply these strategies to other groups of recipients. This could allow us to derive general principles for achieving efficient interspecies conjugation.
4. Conjugation mechanism from the RP4 plasmid (IncP group) is the most widely used and in general offers the most efficient conjugative transfer thus far. This could simply result from the fact that this mechanism has historically been the most studied and therefore well optimized compared to other systems. A few selected studies in our review provide alternatives such as R388 (IncW group) and pCTX-M3 (IncM group). Notably, pIP501, a conjugative plasmid from the Inc18 group has also demonstrated the ability to mediate bidirectional transfer between gram-positive and gram-negative bacteria. Its natural capability to cross the gram barrier makes it a promising candidate for future development as a broad-host-range delivery system. Future work should aim to optimize these systems further and test their capability across a wider range of recipient cells. Having many options for delivery mechanisms would help fill the recipient niches where the traditional RP4 systems cannot access.
5. Surprisingly, none of the selected studies attempted to investigate molecular mechanisms behind observed differences in conjugation efficiencies. One minor exception is the study by Liu et al (2019) that looked into the association between level of ROS, mRNA expression and conjugation efficiency [42]. Conjugation mechanisms thus far have been explored mostly in the context of intraspecies conjugation. It is not known, for example, how regulations of Tra genes and the formation of conjugative pili structures in the donor cells differ as they interact with different recipient cell types. Nor that we know detail kinetic differences of DNA passing into and establishing inside different recipient cells. Answering these questions in the future work would allow us to build models that explain and predict conjugation efficiency across recipient cells type as well as guiding our design of a universal DNA delivery platform.

## Materials and Methods

### Search strategy

A systematic search of the PubMed, Crossref and Web of Science database (up to July 2025), with the following search terms: (conjugation OR conjugative OR conjugal OR mobilizable OR mobilizable) AND ((gram positive) OR gram-positive OR *Bacillus* OR *Bacilli* OR *Lactobacillus* OR *Lactobacilli* OR *Actinomyces* OR *Streptomyces* OR *Corynebacterium*) AND (plasmid* OR vector*). Additionally, we added three studies related to our research objectives [44-46]. These studies were not found in the initial database search but were considered valuable additions to our review.

### Study selection

We screen the list of studies we got from the search above based on title, abstract and full text. The screening was conducted by PS and SC. A study was included if: 1) it provided experimental evidence of plasmid conjugation to at least one gram-positive bacteria, 2) conjugation efficiency was reported. A study was excluded if: 1) no full text was available in English, 2) conjugation efficiency was not reported.

### Data extraction

Data extraction was carried out manually by PS and SC. Both reviewers collaboratively extracted data. No automation tools were used. Study authors were not contacted for additional data. The primary outcome of this review is conjugation efficiency, measured as transconjugant per recipient. Each “study” may consist of multiple conjugation setups, each with different combination of conjugation parameters such as donor strain, recipients’ strains, conjugation machinery, transferred plasmid, cell density, media conditions, etc. Each combination provides a single “measurement” value for our analysis. For repeated experiments, we only take the average conjugation efficiency. The data was organized in a spreadsheet, using a format adapted from Alderliesten et al (2020)[18]. Each row corresponded to data derived from a single measurement, while individual columns provided detailed information about the study, the experimental setup, and the measured conjugation efficiency values. Our data spreadsheet differed from that of Alderliesten et al (2020) in several ways. First, we combined data from both liquid and filter mating experiments into a single spreadsheet, with an additional column specifying the experimental medium as either liquid or solid agar. Second, we used transconjugant per recipient as a metric for conjugation efficiency, as opposed to transconjugant per donor. In cases where the transconjugant per recipient value was not explicitly provided in the study, we estimated it based on the transconjugant per donor, assuming a consistent donor-to-recipient ratio throughout the experiment. If the donor-to-recipient ratio was not available, we assumed it to be 1:1. We did not include measurement data from negative control, i.e., conjugation experiment where conjugation is theoretically impossible, for example, an experiment where OriT or some essential genes in conjugation process was clearly missing. The minimum conjugation efficiency is set to 1E-12 whenever a study reports zero conjugation efficiency or ‘below detection limit’ efficiency. This minimal value helps avoid data visualization problem in log-scale. This value was arbitrary chosen as it is clearly (∼ 3 orders of magnitude) below the lowest measurable values in all of our selected studies. We assessed the potential risk of bias by reporting study size distributions (Fig 1A) and analyzing possible study selection bias (Fig S1). Differences in experimental conditions, bacterial strains, and conjugation setups across studies were considered as potential sources of bias. No formal bias assessment tool was used, but we discussed these limitations in the interpretation of our findings.

### Data analysis

The data from our organized spreadsheet in CSV format was imported to Python as a data frame. Data synthesis involved pooling conjugation efficiency measurements from all selected studies. To assess heterogeneity, subgroup analyses were conducted based on bacterial recipient taxa, conjugation machineries, and experimental conditions. We used log-transformed efficiency distributions to account for variations in measurement scales. Sensitivity analyses were performed by separately analyzing studies that reported both *E. coli* to *E. coli* and *E. coli* to gram-positive conjugation, allowing for direct within-study comparisons. We used Python libraries such as matplotlib.pyplot and seaborn to create various graphical representations, including charts and graphs. We conducted statistical testing using the statsmodels.stats.multicomp module in Python, which facilitated statistical analyses like ANOVA and Tukey’s Honestly Significant Difference (Tukey HSD) tests. The primary effect measure in this study is conjugation efficiency, expressed as transconjugants per recipient. Comparisons across studies were made using mean values and log-transformed efficiency distributions to account for variations in experimental conditions. Statistical significance was assessed using ANOVA and Tukey’s Honest Significant Difference (HSD) tests, as implemented in the statsmodels.stats.multicomp module in Python. Generative AI (ChatGPT) was used to guide python programming and language editing. The Excel file detailing selected studies and extract data as well as iPython Notebook for data analysis were provided in the supplementary of this manuscript. Potential reporting bias was assessed by analyzing study selection patterns and the distribution of reported conjugation efficiencies (Fig S1). Certainty in the findings was assessed by performing sensitivity analyses, comparing results across different subsets of studies. Specifically, we examined whether trends remained consistent when restricting the analysis to studies that reported both *E. coli* to *E. coli* and *E. coli* to gram-positive conjugation. No formal certainty assessment framework (e.g., GRADE) was applied, but robustness of conclusions was evaluated based on consistency across study subsets. No formal statistical test (e.g., funnel plot analysis) was performed to quantify publication bias.

## Supporting information

supplementary result

## Acknowledgement

This study is financially supported by the Air Force Office of Scientific Research, USA, under award number FA2386-23-1-4017, Ministry of Higher Education, Science, Research and Innovation, Thailand, under award number RGNS 63-131, Naresuan University Fundamental Fund 2025 award number 68A107000095. We would also like to thank Faculty of Medical Science, Naresuan University for supporting all facilities. The funding sources had no role in study design, data collection, analysis, or manuscript preparation. The views expressed in this article are those of the authors and do not necessarily reflect the views of the funders.

## Competing Interest

The authors declare that they have no competing interests related to this study.

## Notes

### Competing Interest Statement

The authors have declared no competing interest.

### Summary of Updates

Study search was revised to a more recent date; more studies were included in the analysis

