## supplementary result for "Conjugative Transfer from *Escherichia coli* to Gram-positive Bacteria: A Systematic Review and Meta-Analysis"

1. CSV file of extracted data
   https://docs.google.com/spreadsheets/d/1TLSP4jS6rDjyhUGP6KpsYPpBwvKU3BZI/edit?usp=sharing&ouid=115926259707738595890&rtpof=true&sd=true

2. Python Notebook (on Google Colab) for data visualization and analysis
   https://colab.research.google.com/drive/1DmCKahcEy4F6fAsf2UEeVKlKv2K67CUz?usp=sharing

3. Conjugation efficiency in different recipient groups

### Table S1: Comparing results of statistical tests in using different data sets

| Comparison | | All Data | Internal control | IncP only | Big Studies | Small Studies |
|---|---|---|---|---|---|---|
| Gr 1 (*E.coli*) | Gr 2 (Non *E.coli* G-) | ns | ns | ns | - | ns |
| Gr 1 (*E.coli*) | Gr 3.1 (*Streptomyces*) | *** | *** | * | *** | ** |
| Gr 1 (*E.coli*) | Gr 3.2 (Bacillaceae) | *** | ** | *** | *** | *** |
| Gr 1 (*E.coli*) | Gr 3.3 (Lactobacillaceae) | *** | *** | *** | *** | - |
| Gr 1 (*E.coli*) | Gr 3.4 (Other G+) | *** | *** | *** | *** | *** |
| Gr 2 (Non *E.coli* G-) | Gr 3.1 (*Streptomyces*) | ns | ** | ns | - | ns |
| Gr 2 (Non *E.coli* G-) | Gr 3.2 (Bacillaceae) | *** | ns | ns | - | *** |
| Gr 2 (Non *E.coli* G-) | Gr 3.3 (Lactobacillaceae) | *** | *** | * | - | - |
| Gr 2 (Non *E.coli* G-) | Gr 3.4 (Other G+) | *** | ** | ns | - | *** |
| Gr 3.1 (*Streptomyces*) | Gr 3.2 (Bacillaceae) | *** | ns | *** | ns | *** |
| Gr 3.1 (*Streptomyces*) | Gr 3.3 (Lactobacillaceae) | *** | ns | ** | *** | - |
| Gr 3.1 (*Streptomyces*) | Gr 3.4 (Other G+) | ns | ns | *** | ns | *** |
| Gr 3.2 (Bacillaceae) | Gr 3.3 (Lactobacillaceae) | * | ** | ns | * | - |
| Gr 3.2 (Bacillaceae) | Gr 3.4 (Other G+) | ns | ns | ns | ns | ns |
| Gr 3.3 (Lactobacillaceae) | Gr 3.4 (Other G+) | * | ** | ns | *** | - |

**All data:** full data set from 41 studies with a total of 424 measurements (same as in Fig 3). Note that 380 measurements are from *E.coli*-to-gram positive conjugation; the rest are from *E.coli*-to-gram negative conjugation.
**Internal control:** data only from 10 studies that have both *E.coli*-to-*E.coli* and *E.coli*-to-gram positive conjugation efficiency measures; these studies provide 193 measurements in total.
**IncP only**: data only from experiments that employed conjugation mechanisms from Inc P plasmid groups; there are 346 measurements in total.
**Big studies:** data only from 10 studies that have more than 10 measurements; these studies alone provide 290 measurements in total.
**Small studies:** data only from 31 studies that have no more than 10 measurements; these studies alone provide 134 measurements in total.

p-value from post hoc analysis: ns $P > 0.05$; * $P \leq 0.05$; ** $P \leq 0.01$; *** $P \leq 0.001$, - no data

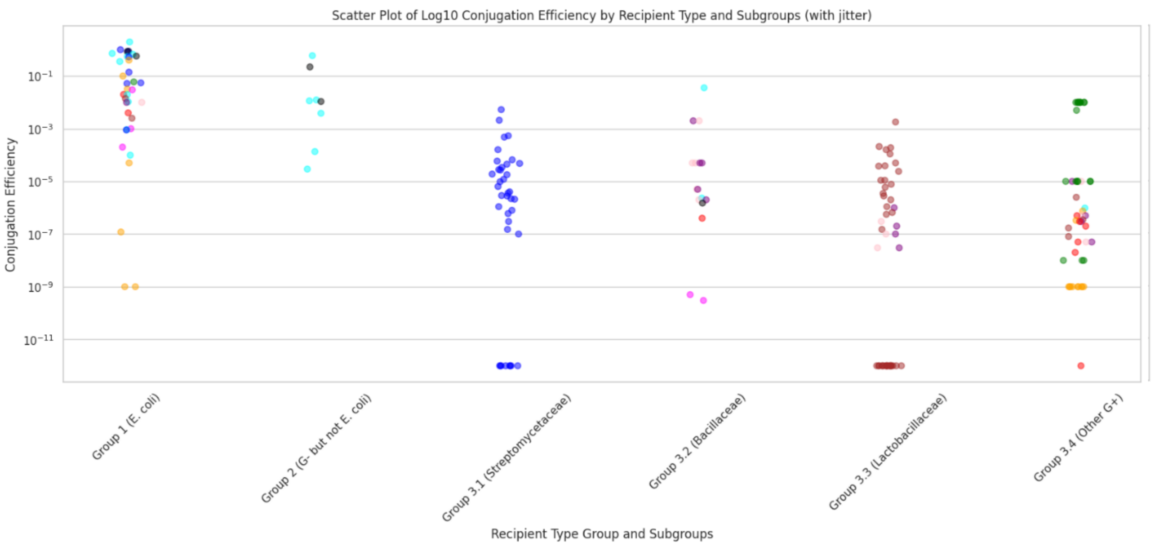


data groups are statistically different

p-value = 3.9356744355214955e-18

Multiple Comparison of Means - Tukey HSD, FWER=0.05

| group1 | group2 | meandiff | p-adj | lower | upper | reject |
|---|---|---|---|---|---|---|
| Group 1 (E. coli) | Group 2 (G- but not E. coli) | -0.1687 | 1.0 | -3.3229 | 2.9855 | False |
| Group 1 (E. coli) | Group 3.1 (Streptomycetaceae) | -4.4253 | 0.0 | -6.3145 | -2.5361 | True |
| Group 1 (E. coli) | Group 3.2 (Bacillaceae) | -3.1502 | 0.0033 | -5.5749 | -0.7255 | True |
| Group 1 (E. coli) | Group 3.3 (Lactobacillaceae) | -6.1943 | 0.0 | -7.9816 | -4.407 | True |
| Group 1 (E. coli) | Group 3.4 (Other G+) | -4.0019 | 0.0 | -5.7733 | -2.2305 | True |
| Group 2 (G- but not E. coli) | Group 3.1 (Streptomycetaceae) | -4.2566 | 0.0019 | -7.4031 | -1.1102 | True |
| Group 2 (G- but not E. coli) | Group 3.2 (Bacillaceae) | -2.9815 | 0.1426 | -6.4758 | 0.5128 | False |
| Group 2 (G- but not E. coli) | Group 3.3 (Lactobacillaceae) | -6.0256 | 0.0 | -9.112 | -2.9393 | True |
| Group 2 (G- but not E. coli) | Group 3.4 (Other G+) | -3.8332 | 0.0056 | -6.9104 | -0.756 | True |
| Group 3.1 (Streptomycetaceae) | Group 3.2 (Bacillaceae) | 1.2751 | 0.6514 | -1.1394 | 3.6897 | False |
| Group 3.1 (Streptomycetaceae) | Group 3.3 (Lactobacillaceae) | -1.769 | 0.051 | -3.5426 | 0.0046 | False |
| Group 3.1 (Streptomycetaceae) | Group 3.4 (Other G+) | 0.4234 | 0.9825 | -1.3341 | 2.181 | False |
| Group 3.2 (Bacillaceae) | Group 3.3 (Lactobacillaceae) | -3.0441 | 0.0031 | -5.3799 | -0.7084 | True |
| Group 3.2 (Bacillaceae) | Group 3.4 (Other G+) | -0.8517 | 0.8982 | -3.1753 | 1.4719 | False |
| Group 3.3 (Lactobacillaceae) | Group 3.4 (Other G+) | 2.1924 | 0.0024 | 0.5448 | 3.84 | True |

**Fig. S1 Conjugation efficiency from *E.coli* donor to various groups of recipient bacteria, using only data set from studies with both *E.coli*-to-*E.coli* and *E.coli*-to-gram positive conjugation.** Different colors denote studies where data points come from: ID: color = {46: 'red', 39: 'green', 23: 'blue', 55: 'purple', 32: 'orange', 56: 'pink',24: ‘cyan’, 54: 'magenta', 31: 'brown', 18: 'black',}

A

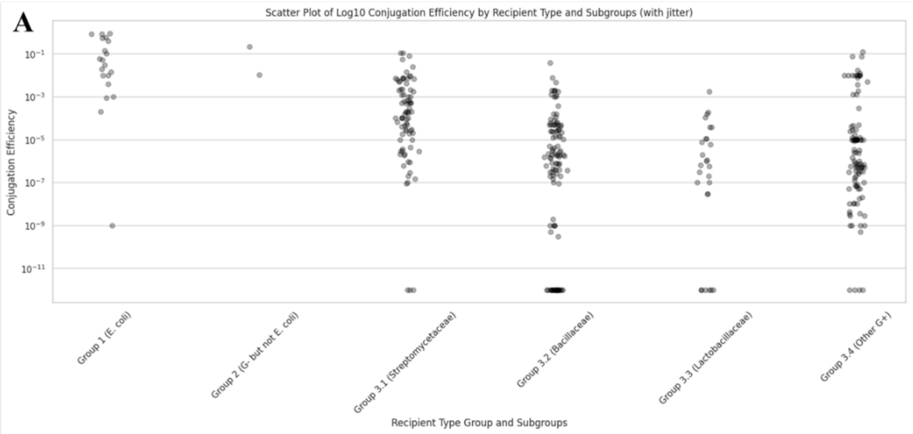


data groups are statistically different
p-value = 3.698480799808228e-17

Multiple Comparison of Means - Tukey HSD, FWER=0.05

| group1 | group2 | meandiff | p-adj | lower | upper | reject |
|---|---|---|---|---|---|---|
| Group 1 (E. coli) | Group 2 (G- but not E. coli) | 0.4734 | 0.9999 | -5.179 | 6.1258 | False |
| Group 1 (E. coli) | Group 3.1 (Streptomycetaceae) | -2.3116 | 0.0074 | -4.2146 | -0.4085 | True |
| Group 1 (E. coli) | Group 3.2 (Bacillaceae) | -4.917 | 0.0 | -6.7672 | -3.0668 | True |
| Group 1 (E. coli) | Group 3.3 (Lactobacillaceae) | -5.1655 | 0.0 | -7.4141 | -2.917 | True |
| Group 1 (E. coli) | Group 3.4 (Other G+) | -3.9031 | 0.0 | -5.7641 | -2.0422 | True |
| Group 2 (G- but not E. coli) | Group 3.1 (Streptomycetaceae) | -2.785 | 0.688 | -8.2405 | 2.6705 | False |
| Group 2 (G- but not E. coli) | Group 3.2 (Bacillaceae) | -5.3904 | 0.0535 | -10.8277 | 0.0468 | False |
| Group 2 (G- but not E. coli) | Group 3.3 (Lactobacillaceae) | -5.639 | 0.0463 | -11.2244 | -0.0536 | True |
| Group 2 (G- but not E. coli) | Group 3.4 (Other G+) | -4.3766 | 0.1947 | -9.8175 | 1.0644 | False |
| Group 3.1 (Streptomycetaceae) | Group 3.2 (Bacillaceae) | -2.6054 | 0.0 | -3.7171 | -1.4938 | True |
| Group 3.1 (Streptomycetaceae) | Group 3.3 (Lactobacillaceae) | -2.854 | 0.0 | -4.5477 | -1.1603 | True |
| Group 3.1 (Streptomycetaceae) | Group 3.4 (Other G+) | -1.5916 | 0.0009 | -2.7211 | -0.4621 | True |
| Group 3.2 (Bacillaceae) | Group 3.3 (Lactobacillaceae) | -0.2485 | 0.998 | -1.8826 | 1.3855 | False |
| Group 3.2 (Bacillaceae) | Group 3.4 (Other G+) | 1.0139 | 0.0599 | -0.024 | 2.0517 | False |
| Group 3.3 (Lactobacillaceae) | Group 3.4 (Other G+) | 1.2624 | 0.2414 | -0.3838 | 2.9086 | False |

B

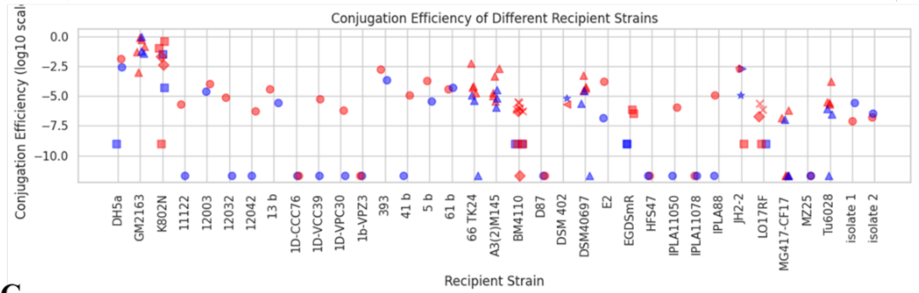


C

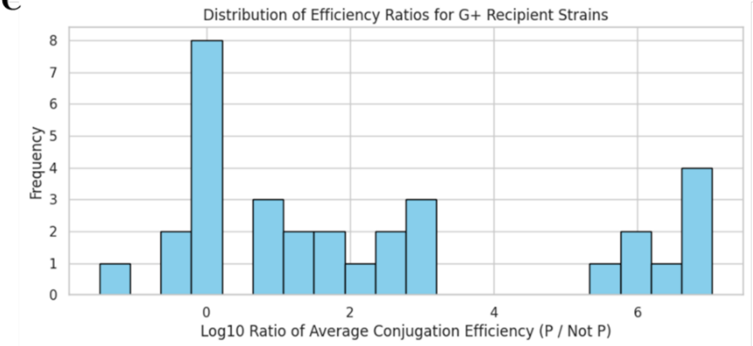


**Fig. S2 Conjugation efficiency and conjugation machinery.** (A) Conjugation efficiency between *E.coli* donor and various groups of recipient bacteria, using only data set from experiments that use conjugation machinery from IncP plasmid group. (B) Conjugation efficiencies across different recipient strains when using conjugation machinery from Inc P (red) or other plasmid groups (blue). The first two strains are *E.coli*; the last two strains are *S. epidermidis* from [31] (C) Distribution of log10 ratio between conjugation efficiency in different recipient strains when Inc P or not Inc P conjugation machineries were used.

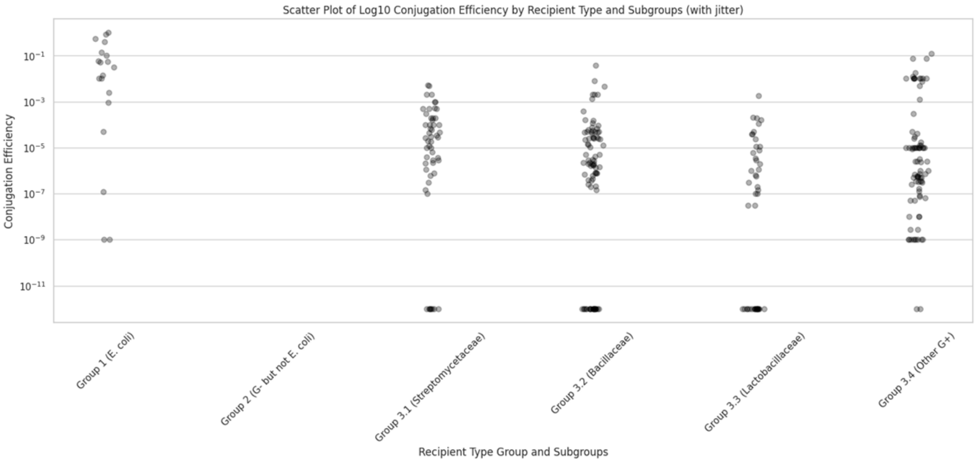


data groups are statistically different
p-value = 1.1210069340952457e-10

Multiple Comparison of Means - Tukey HSD, FWER=0.05

| group1 | group2 | meandiff | p-adj | lower | upper | reject |
|---|---|---|---|---|---|---|
| Group 1 (E. coli) | Group 3.1 (Streptomycetaceae) | -3.0324 | 0.0017 | -5.2314 | -0.8334 | True |
| Group 1 (E. coli) | Group 3.2 (Bacillaceae) | -4.1396 | 0.0 | -6.2242 | -2.0549 | True |
| Group 1 (E. coli) | Group 3.3 (Lactobacillaceae) | -5.6577 | 0.0 | -7.8934 | -3.4219 | True |
| Group 1 (E. coli) | Group 3.4 (Other G+) | -3.0334 | 0.0008 | -5.1203 | -0.9465 | True |
| Group 3.1 (Streptomycetaceae) | Group 3.2 (Bacillaceae) | -1.1072 | 0.2159 | -2.5433 | 0.3289 | False |
| Group 3.1 (Streptomycetaceae) | Group 3.3 (Lactobacillaceae) | -2.6253 | 0.0002 | -4.273 | -0.9776 | True |
| Group 3.1 (Streptomycetaceae) | Group 3.4 (Other G+) | -0.001 | 1.0 | -1.4404 | 1.4383 | False |
| Group 3.2 (Bacillaceae) | Group 3.3 (Lactobacillaceae) | -1.5181 | 0.0438 | -3.0099 | -0.0264 | True |
| Group 3.2 (Bacillaceae) | Group 3.4 (Other G+) | 1.1062 | 0.1144 | -0.1516 | 2.364 | False |
| Group 3.3 (Lactobacillaceae) | Group 3.4 (Other G+) | 2.6243 | 0.0 | 1.1294 | 4.1192 | True |

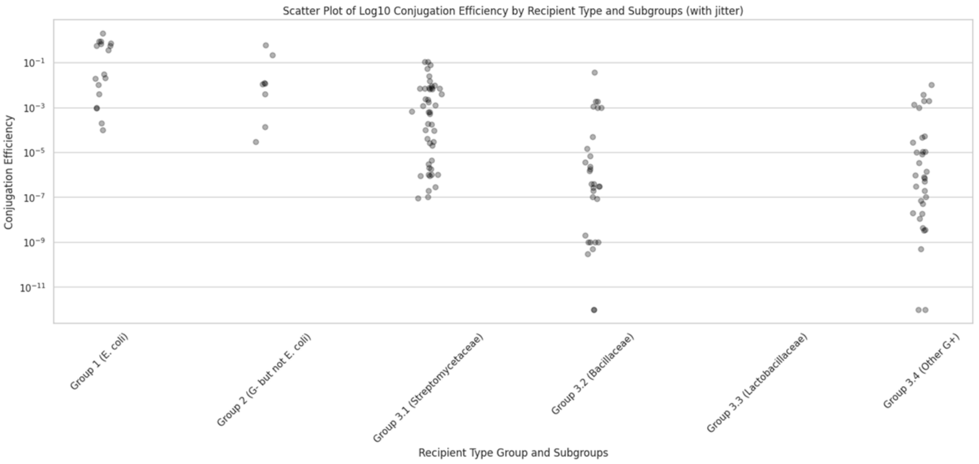


data groups are statistically different
p-value = 1.7414520297028193e-14

Multiple Comparison of Means - Tukey HSD, FWER=0.05

| group1 | group2 | meandiff | p-adj | lower | upper | reject |
|---|---|---|---|---|---|---|
| Group 1 (E. coli) | Group 2 (G- but not E. coli) | -0.7685 | 0.9209 | -3.3307 | 1.7937 | False |
| Group 1 (E. coli) | Group 3.1 (Streptomycetaceae) | -2.2606 | 0.003 | -3.9568 | -0.5644 | True |
| Group 1 (E. coli) | Group 3.2 (Bacillaceae) | -5.0302 | 0.0 | -6.8443 | -3.216 | True |
| Group 1 (E. coli) | Group 3.4 (Other G+) | -4.6843 | 0.0 | -6.4684 | -2.9003 | True |
| Group 2 (G- but not E. coli) | Group 3.1 (Streptomycetaceae) | -1.4921 | 0.3761 | -3.7813 | 0.7971 | False |
| Group 2 (G- but not E. coli) | Group 3.2 (Bacillaceae) | -4.2617 | 0.0 | -6.6396 | -1.8838 | True |
| Group 2 (G- but not E. coli) | Group 3.4 (Other G+) | -3.9159 | 0.0001 | -6.2709 | -1.5608 | True |
| Group 3.1 (Streptomycetaceae) | Group 3.2 (Bacillaceae) | -2.7696 | 0.0 | -4.172 | -1.3672 | True |
| Group 3.1 (Streptomycetaceae) | Group 3.4 (Other G+) | -2.4237 | 0.0 | -3.787 | -1.0605 | True |
| Group 3.2 (Bacillaceae) | Group 3.4 (Other G+) | 0.3458 | 0.9691 | -1.1617 | 1.8534 | False |

**Fig. S3 Conjugation efficiency from *E.coli* donor to various groups of recipient bacteria, using only data set from studies with more than 10 measurements (top) or no more than 10 measurements (bottom).**

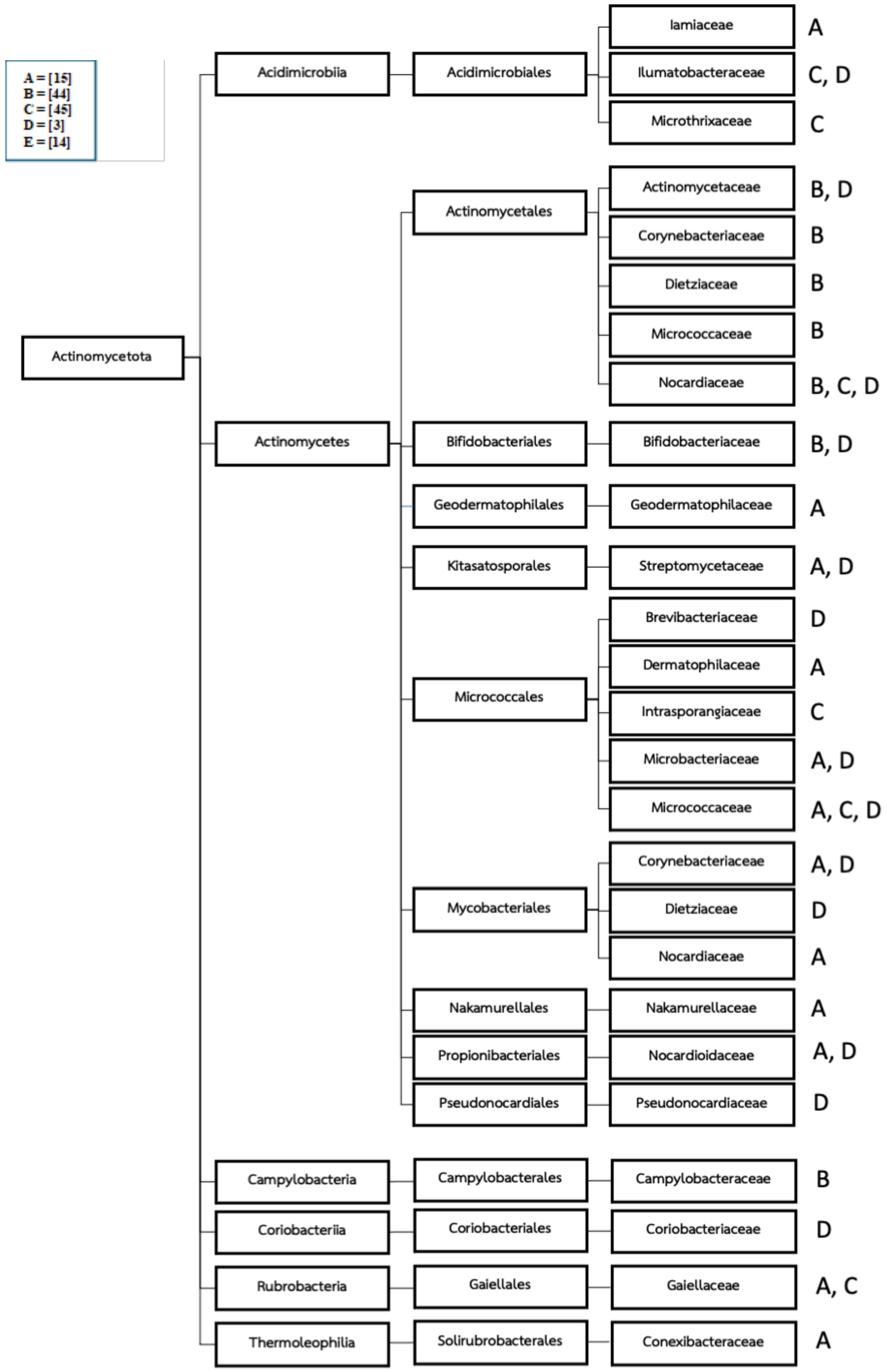
A = [15]
B = [44]
C = [45]
D = [3]
E = [14]
Actinomycetota
Acidimicrobiia
Acidimicrobiales
Iamiaceae
A
Ilumatobacteraceae
C, D
Microthrixaceae
C
Actinomycetes
Actinomycetales
Actinomycetaceae
B, D
Corynebacteriaceae
B
Dietziaceae
B
Micrococcaceae
B
Nocardiaceae
B, C, D
Bifidobacteriales
Bifidobacteriaceae
B, D
Geodermatophilales
Geodermatophilaceae
A
Kitasatosporales
Streptomycetaceae
A, D
Micrococcales
Brevibacteriaceae
D
Dermatophilaceae
A
Intrasporangiaceae
C
Microbacteriaceae
A, D
Micrococcaceae
A, C, D
Mycobacteriales
Corynebacteriaceae
A, D
Dietziaceae
D
Nocardiaceae
A
Nakamurellales
Nakamurellaceae
A
Propionibacteriales
Nocardioidaceae
A, D
Pseudonocardiales
Pseudonocardiaceae
D
Campylobacteria
Campylobacterales
Campylobacteraceae
B
Coriobacteriia
Coriobacteriales
Coriobacteriaceae
D
Rubrobacteria
Gaiellales
Gaiellaceae
A, C
Thermoleophilia
Solirubrobacterales
Conexibacteraceae
A

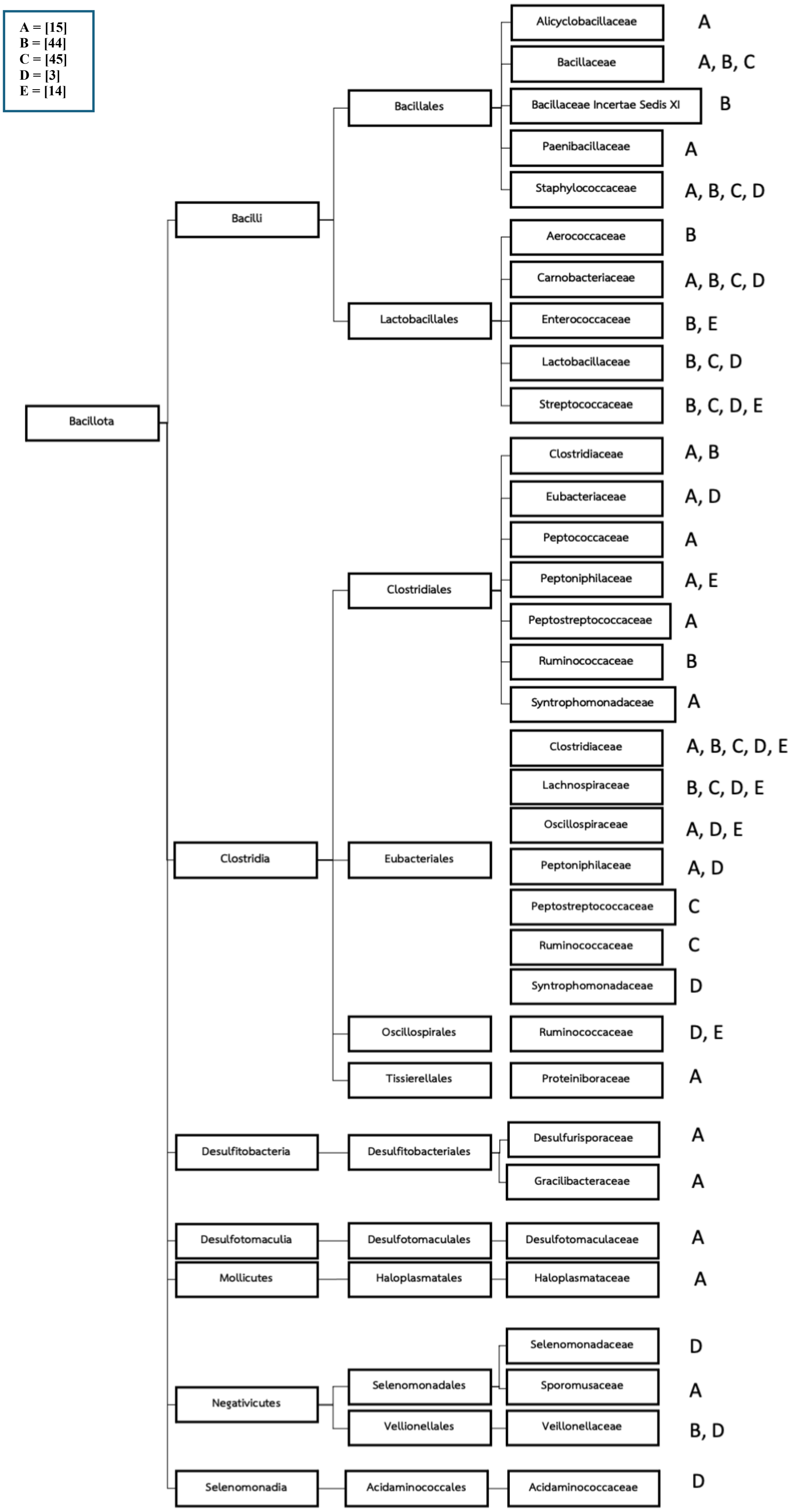


**Fig. S4 Gram+ bacteria families that can receive plasmid from Gram- bacteria (*E.coli* or *P.putida*), according to *in situ* conjugation experiment.**
